# Differential endopeptidase requirements during adaptation to changing growth conditions in *Vibrio cholerae*

**DOI:** 10.1101/2025.01.28.635349

**Authors:** Kelly Rosch, Samantha Lei, Jennifer Zheng, Tobias Dörr

## Abstract

The bacterial cell wall is a covalently linked meshwork of peptidoglycan (PG) that establishes cell shape and prevents osmotic lysis. This structure must be flexible enough to accommodate transenvelope protein complexes, but strong enough to withstand high intracellular pressure. In order to elongate and divide, cells must remodel the cell wall through the concerted action of PG synthesis and degradation. Endopeptidases, a class of PG degrading enzymes, facilitate cell growth by hydrolyzing PG crosslinks. *Vibrio cholerae* encodes several functionally redundant endopeptidases, two of which are nearly identical: ShyA and ShyC. To investigate differential roles of these enzymes, we assessed growth and morphology of ShyA and ShyC mutants. We found that ShyA, but not ShyC, is required for normal adaptation to low osmolarity medium. Cells lacking ShyA exhibited longer lag phase and aberrant morphology during adaptation, and reduced survival in the presence of a beta-lactam antibiotic. Lastly, our experiments revealed that cells lacking ShyA’s LysM domain exhibited more severe defects than cells lacking ShyA altogether, implicating the LysM domain in proper regulation of ShyA activity.

## Introduction

Nearly all bacteria contain a peptidoglycan (PG) cell wall, a covalently linked meshwork that establishes cell shape and protects the cell from osmotic lysis. PG consists of glycan strands, comprising polymerized alternating N-acetylmuramic acid and N-acetylglucosamine sugars, which are connected by peptide crosslinks [1-2]. In Gram-negative bacteria, the cell wall is a thin layer of PG enclosed between the inner and outer membrane in a space called the periplasm [3]. This thin PG layer must be malleable enough to accommodate insertion of transenvelope protein complexes [4], while also being strong enough to withstand intracellular pressure roughly equivalent to that of a car tire [2, 5-6]. In order to elongate and divide, cells must remodel PG without losing structural integrity. Cell wall remodeling requires the activity of PG synthases, which insert new PG material into the existing cell wall [7], and autolysins, a divergent group of enzymes that mediate cell wall turnover and create holes in the cell wall for new PG to be inserted [4, 8-9]. The activity of cell wall synthases and hydrolases is presumably tightly regulated because an imbalance of either activity can lead to severe growth defects or lysis [10]. Indeed, the activity of autolysins is a major contributor to the antibacterial mechanism of the beta-lactam antibiotics, which are among the most widely-prescribed antibiotics worldwide. Beta-lactams inhibit cell wall synthases, resulting in autolysin-mediated cell wall degradation, which causes either cell death and lysis, or inhibition of cell division [11]. However, the mechanism by which this balance is maintained remains unclear.

Bacteria frequently encounter changing environmental conditions during their life cycles, which challenge the integrity of the cell wall and can disrupt the balance between these opposing classes of cell wall remodeling enzymes. *Vibrio cholerae*, the model organism in this study, lives in brackish water, pond water, and in the human gut, where it causes cholera disease [12]. During its life cycle, *V. cholerae* must adapt to changes in osmotic conditions and accommodate the resulting fluctuations in turgor pressure. As cells transition between different environments, substantial PG remodeling is likely required, further challenging the balance among PG remodeling enzymes.

Autolysins have been shown to play a role during environmental adaptation. Indeed, multiple carboxypeptidases and lytic transglycosylases have been shown to respond to changes in pH or outer membrane stress [13]. In *E. coli*, the endopeptidases MepS and MepM are only required under certain nutrient availability conditions [14]. In *V. cholerae*, the endopeptidase ShyB is expressed only during zinc starvation [15]. Additionally, endopeptidases may play a role in cell envelope modifications during mechanical stress; in *V. cholerae*, mechanical perturbations of the cell wall activate the VxrAB cell wall stress response system in a ShyA-dependent manner [16].

Endopeptidases are relatively uncharacterized compared to PG synthases, partially due to their functional redundancy, which makes simple genotype-phenotype dissection challenging. *V. cholerae* encodes nine endopeptidases, many of which remain functionally uncharacterized. Two of these endopeptidases are expressed during normal growth: ShyA and ShyC, which are homologous to the *E. coli* D,D-endopeptidase MepM [8]. ShyA and ShyC are nearly identical, both containing an M23 metalloendopeptidase domain [17] and a LysM carbohydrate binding domain [18]. ShyA is soluble in the periplasm while ShyC is tethered to the inner membrane. ShyA and ShyC are conditionally essential [9]. While both endopeptidases can fulfill normal growth functions in standard laboratory media, the reason for this apparent functional redundancy is unknown.

Here, we assessed the roles of *V. cholerae* endopeptidases ShyA and ShyC during adaptation to low osmolarity. We found that ShyA, but not ShyC, is required for normal adaptation to salt-free medium, indicating different roles for these nearly identical enzymes.

## Results

*Vibrio cholerae* encodes two principal endopeptidases (EPs), ShyA and ShyC, which are constitutively expressed during normal growth. ShyA is soluble in the periplasm, and ShyC is tethered to the inner membrane. ShyA and ShyC are collectively essential, but individually redundant, like most autolysins in most bacteria. The reasons for functional redundancy of autolysins are poorly understood. We hypothesized that endopeptidases may play a role in adjusting PG cleavage during changes in turgor pressure. We thus investigated the requirement for ShyA and ShyC to ensure proper growth and morphogenesis during adaptation to low osmolarity (which results in increased turgor). To this end, we incubated Δ*shyA* and Δ*shyC* mutants in a standard laboratory growth medium with (LB) or without (SF-LB) addition of 180 mM NaCl, and observed cell morphology (Figure 1A). While neither mutant had a morphology defect in LB, cells lacking ShyA exhibited a morphology defect in SF-LB, and often formed phase-light blebs (Figure 1A). In contrast, cells lacking ShyC exhibited normal morphology in SF-LB (Figure 1A), indicating differential roles for these endopeptidases in a low osmolarity environment. We next used the PG label BADA and the membrane stain FM4-64 to visualize envelope components. The blebs produced in the *ΔshyA* mutant were stained with FM4-64, but contained qualitatively reduced PG labeling compared to the cell body, indicating that this aberrant structure consists primarily of the outer membrane (Figure 1B). These results are consistent with a lack of PG expansion in the *shyA* mutant, which causes an imbalance in the production of cell envelope components; this is also reminiscent of the large outer membrane blebs we previously observed in strains lacking EP activity entirely [9, 19].

**Figure 1.**
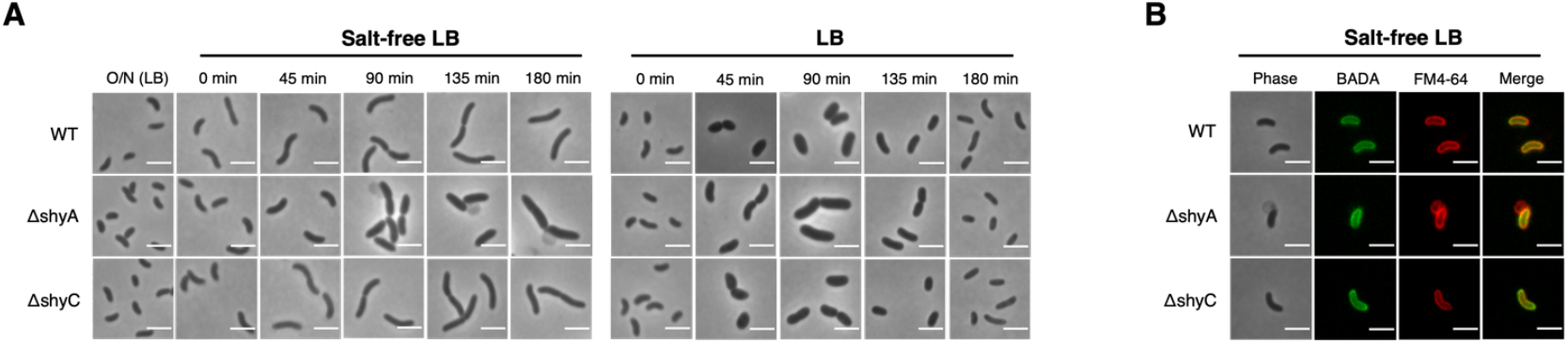
Cells missing ShyA exhibit a morphology defect in low osmolarity medium. **(A)** After overnight incubation, cultures were diluted 1:1000 into LB or SF-LB, incubated at 37C, and imaged on an agarose pad (1.6%) at the indicated timepoints. Scale bar = 3 microns. **(B)** BADA and FM4-64 staining after 135 min incubation in SF medium. Scale bar = 3 microns.

We were then curious about the time scale of the *ΔshyA* defect in LB-SF medium – does this defect resolve over time? To investigate this, we plated cells lacking ShyA, and cells expressing ShyA *in trans*, on LB and SF-LB and imaged the plates after 12 hours and 36 hours. Consistent with the observed morphology defects, cells lacking ShyA exhibited a slight but reproducible growth defect on SF-LB but not on LB (Figure 2A). This growth defect on low osmolarity medium became less pronounced after extended incubation, indicating that damage primarily occurs during the initial stages of adaptation to SF medium (Figure 2A). Cells lacking ShyC did not exhibit a growth defect on LB or SF-LB (Supplemental 2A), and overexpression of ShyC could not rescue the *ΔshyA* growth defect on SF-LB (Supplemental 2B), again supporting a more important role for ShyA compared to ShyC in adaptation to low osmolarity.

**Figure 2.**
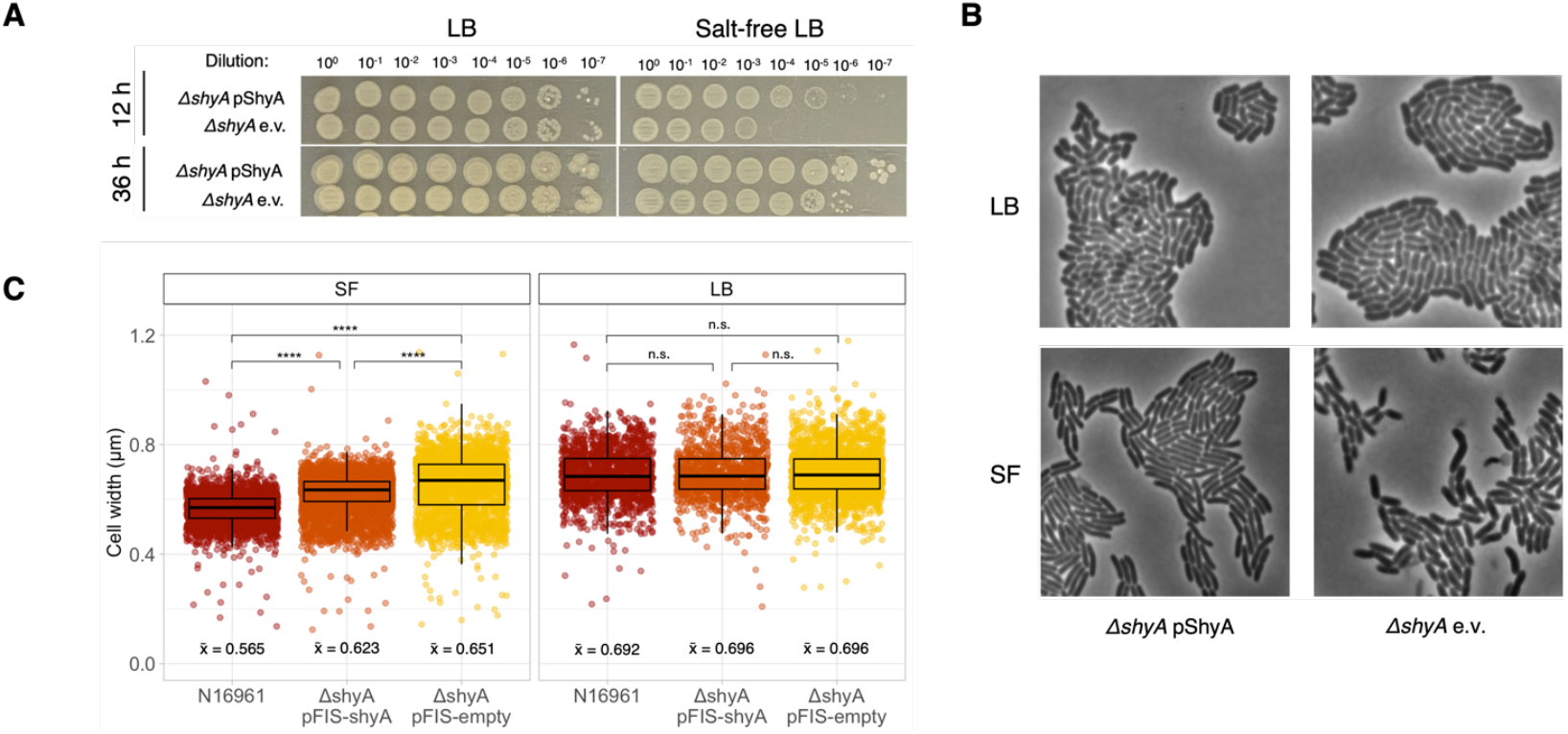
Cells lacking ShyA exhibit transient growth and morphology defects on salt-free medium. **(A)** Spot dilutions of ShyA mutants were incubated at 37C on LB or salt-free LB and imaged after 12 hours and 36 hours. **(B)** ShyA mutants were imaged for 2.5 h using time-lapse microscopy on a 1.6% agarose pad containing LB medium or salt-free LB medium. **(C)** Cell length and width were quantified using SuperSegger; p-values produced by pairwise comparisons using a linear model; n.s. = not significant, p < 0.0001 ****

Next we asked how cell morphology changes over time when cells are grown on a low osmolarity agarose pad. While cells lacking ShyA or expressing ShyA *in trans* formed normal microcolonies on an LB agarose pad, the Δ*shyA* mutant failed to produce normal microcolonies on SF-LB (Figure 2B). Instead, these cells exhibited a similar blebbing phenotype as observed in liquid medium, combined with qualitatively delayed growth (Figure 1A, Figure 1B). On SF medium, cells lacking ShyA were also significantly wider than both WT and cells expressing ShyA *in trans* (Figure 2C), while on LB medium none of the strains varied significantly in width (Figure 2C). On SF-LB, cells expressing ShyA *in trans* exhibited an intermediate cell width between WT and Δ*shyA*, possibly due to a lower ShyA expression level from our inducible promoter compared to WT.

We noticed that cells lacking ShyA formed smaller microcolonies on the SF agarose pad, and thus hypothesized that these cells divided more slowly than their counterparts. To investigate this, we analyzed time lapse images using SuperSegger [20], a software that enables lineage tracking for individual cells. Indeed, on SF medium, cells lacking ShyA took significantly longer to divide (mean = 35.5 mins +/-28.9) than WT cells (17.4 mins +/-15.2), or cells expressing ShyA *in trans* (21.5 mins +/-16.5) (Figure 3A). Conversely, all strains divided normally (12-13 mins) when applied to an agarose pad containing LB medium (Figure 3A).

**Figure 3.**
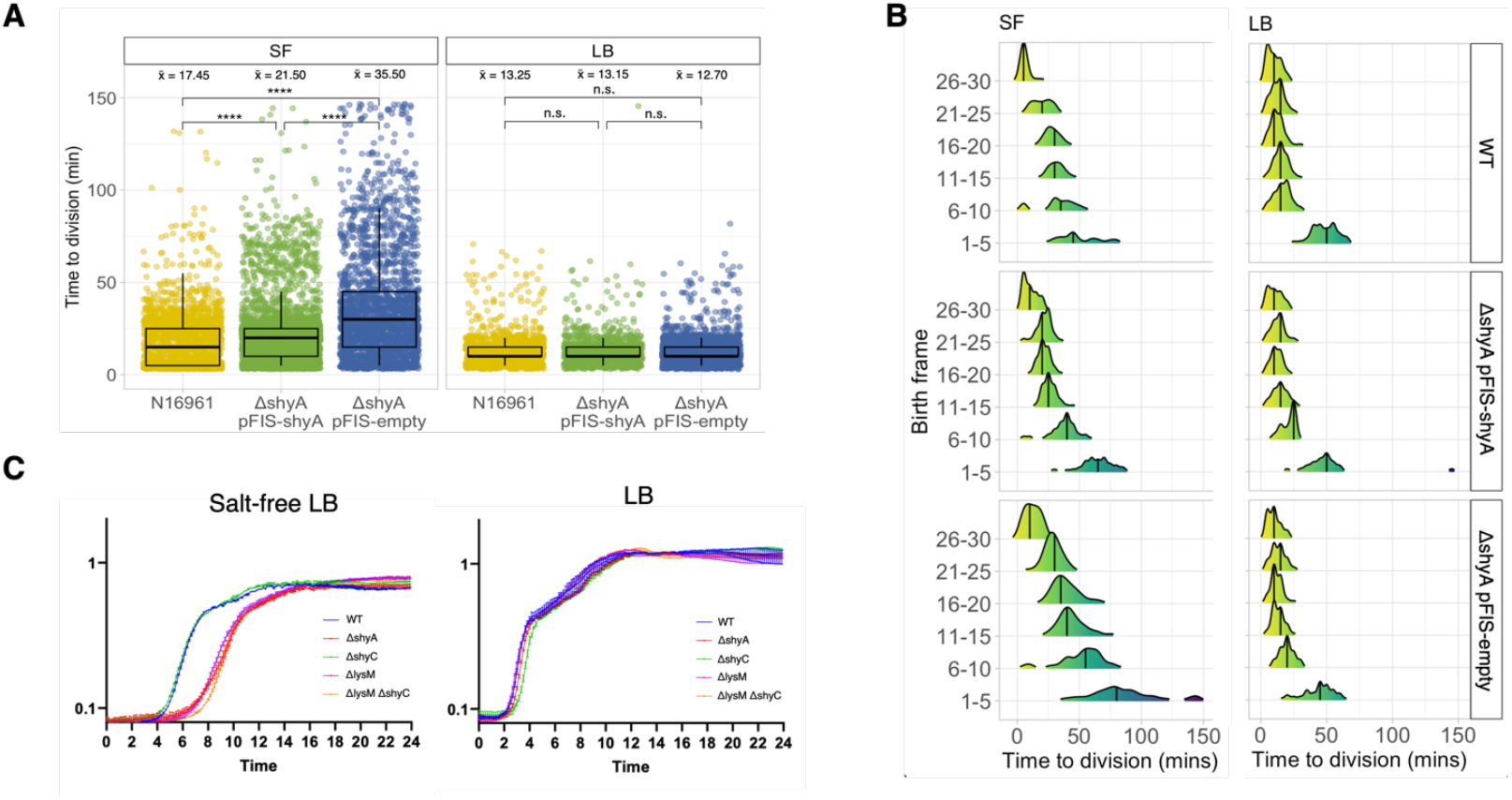
ShyA mutants exhibit increased time to division on low osmolarity medium. **(A)** Time to division for ShyA mutants was quantified using SuperSegger; each point is one cell. **(B)** Time to division binned by birth frame (the frame in which the cell was first detected by SuperSegger); frames taken every 5 min. **(C)** OD600 measurements taken every 10 min during incubation at 37C.

We then sought to distinguish adaptation to low osmolarity from general growth in SF-LB. To investigate this, we binned cells based on the time lapse frame in which they were first detected by SuperSegger (“Birth frame”) to ask whether defects were more pronounced early in the time lapse (adaptation phase defect) or evenly distributed throughout the time lapse (overall growth defect). Indeed, on SF medium the division time for cells lacking ShyA was longest during the early frames of the time lapse, and shortened as the time lapse progressed (Figure 3B), suggesting that ultimately, the Δ*shyA* mutant can adapt to low osmolarity. Conversely, on LB medium, the division time for all strains was slightly elevated during frames 1-5, presumably due to cells exiting lag phase, and then remained short for the duration of the time lapse (Figure 3B). Consistent with the single-cell observations, bulk populations of cells lacking ShyA took longer to adapt and to grow in liquid SF medium than cells expressing ShyA, while in LB cells lacking ShyA had no disadvantage (Figure 3C). Together, these data indicate that the *ΔshyA* growth defect in SF medium is most pronounced during initial adaptation to low osmolarity.

ShyA contains a LysM carbohydrate binding domain that is predicted to bind to the glycan strands in PG [18]. We reasoned that if the LysM domain is involved in PG binding, it might play a crucial role during ShyA-mediated adaptation to low osmolarity. To investigate this, we created a mutant lacking the LysM domain of ShyA (ShyA^ΔlysM^). We were also able to construct a mutant lacking both the LysM domain and ShyC (ShyA^ΔlysM^ *ΔshyC*), demonstrating that ShyA^ΔlysM^ is functional at least during growth in standard laboratory media. We next observed the morphology of these mutants in LB and SF-LB. Like *ΔshyA*, both of these mutants exhibited morphology defects in SF LB, but not in LB (Figure 4A). Both ShyA^ΔlysM^ strains showed bulbous protrusions, but unlike *ΔshyA*, these protrusions were phase dark instead of phase light, indicating that they contained cytoplasm. Cell wall staining indicated that the protrusions contained qualitatively less PG than the cell body, indicating significant local cell wall degradation (Figure 4B). Thus, the LysM domain of ShyA is required for adaptation to low osmolarity, and the ShyA^ΔlysM^ strain suffers from increased PG degradation during transition to lower osmolarity. This may suggest that ShyA’s cell wall binding capability is required for regulating its activity under these conditions – i.e. without LysM, ShyA activity becomes unrestrained in some cells.

**Figure 4.**
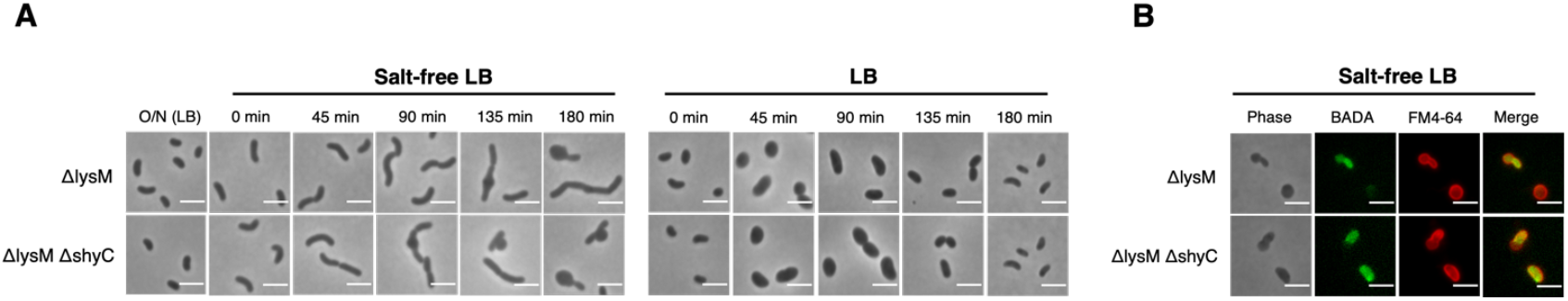
ShyA phenotypes in low osmolarity medium are dependent on the LysM domain. **(A)** After overnight incubation, cultures were diluted 1:1000 into LB or SF-LB, incubated at 37C, and imaged on an agarose pad (1.6%) at the indicated timepoints. Scale bar = 3 microns. **(B)** BADA and FM4-64 staining after 135 min incubation in SF medium. Scale bar = 3 microns.

We next reasoned that the increased lag phase of *ΔshyA* mutants may represent a trade-off with antimicrobial susceptibility. Beta-lactam antibiotics only affect growing cells, and enhanced lag phase is a well-characterized mechanism of antibiotic tolerance [25]. We thus diluted our mutants into LB and SF-LB containing Penicillin G (100 µg/mL, 10x MIC) and measured survival over time by plating for CFU/mL. Consistent with lag-phase-dependent tolerance, WT *V. cholerae* survived significantly better in SF-LB (longer lag phase) than in LB medium (Fig. 5AB). However, contrary to our expectations, cells lacking ShyA exhibited considerably lower survival in the presence of Penicillin G compared to WT or *ΔshyC*, both in LB and SF-LB (Figure 5A). This observation suggests that even though Δ*shyA* cells do not divide during transition into SF-LB, they experience significantly perturbed cell wall turnover, which exacerbates cell wall damage induced by Penicillin G exposure. Indeed, cells lacking only the LysM domain of ShyA (which exhibit more severe cell envelope homeostasis defects based on our single-cell microscopy assays) were even more sensitive to Penicillin G than cells lacking ShyA altogether, indicating that this mutant may have more pronounced structural cell wall defects (Figure 5AB). This trend was exacerbated in LB compared to SF-LB, likely due to the increased growth rate in LB, and thus larger impact of beta-lactam treatment (Figure 5B).

**Figure 5.**
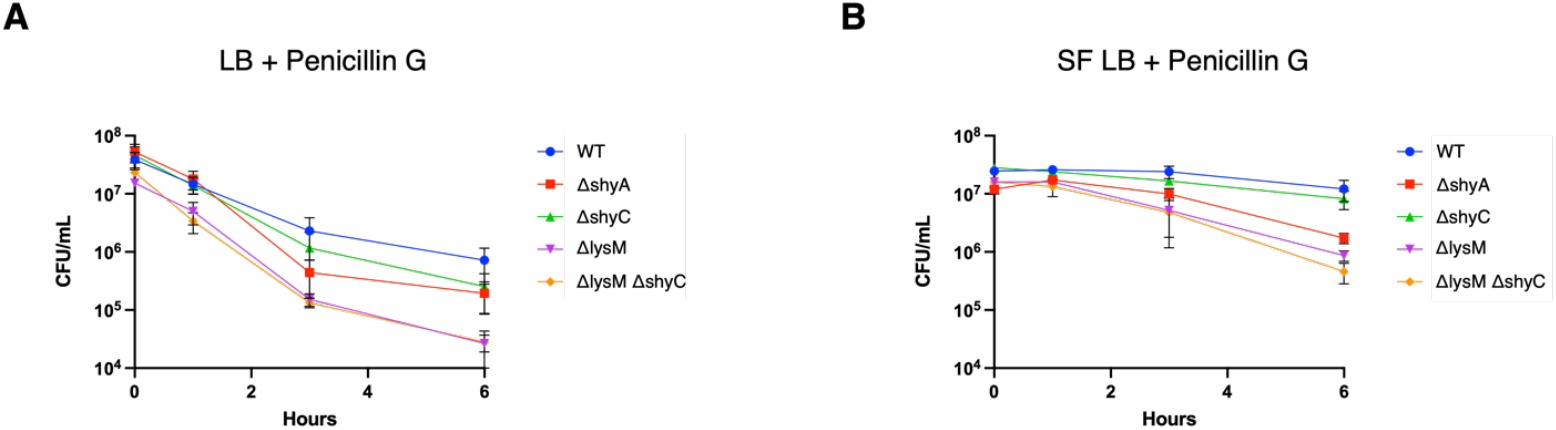
The ShyA LysM domain confers antibiotic tolerance. **(A)-(B)** After overnight incubation, strains were subcultured 1:100 into LB or SF-LB containing 100 ug/mL Penicillin G. Viable cell counts (CFU/mL) were determined at the indicated time points. Error bars represent SEM of three biological replicates.

## Discussion

Here we show that the *V. cholerae* endopeptidase ShyA is important for adaptation to low osmolarity during transition out of stationary phase. While most studies on osmolarity challenges have focused on cytoplasmic solute homeostasis (e.g., cation exchange and compatible solutes [21-22), we explored here the importance of endopeptidase activity in responding to osmotic stress on the cell wall. This is an important finding because it shows that cell wall remodeling may be a crucial and underappreciated component during bacterial life cycles as cells transition into new environments. Interestingly, Δ*shyA* cells adapting to low osmolarity resembled phenotypes observed during general endopeptidase insufficiency (OM blebbing [9, 19]), suggesting that under these conditions, cell wall cleavage is a limiting factor for optimal growth. As a side note, the outer membrane blebbing we observed suggests that *V. cholerae* may not be able to modulate outer membrane biosynthesis relative to cell wall biosynthesis when endopeptidase activity is limited.

Cells encode multiple copies of each class of autolysins, indicating that these redundant genes may play differential roles under different conditions. Our work indicates a novel type of specialization for autolysins: an endopeptidase that specializes in adaptation to low osmolarity. The LysM carbohydrate binding domain in ShyA appears to be critical for ShyA’s function under these conditions. While LysM function in endopeptidases is unknown, this domain is clearly central to ShyA function, as cells lacking the LysM domain of ShyA perform worse than cells lacking ShyA altogether.

Our data also implicate the ShyA LysM domain in antibiotic tolerance. While cells lacking ShyA exhibited a slight sensitivity to Penicillin G in SF-LB, cells lacking the LysM domain of ShyA show exacerbated sensitivity to Penicillin G, especially in LB, where growth rate is increased. This indicates that the endopeptidase ShyA is important for cell wall remodeling during a variety of environmental conditions, from changing osmolarity during the normal life cycle to antibiotic treatment in the clinic.

## Materials and Methods

### Bacterial growth conditions

Liquid cultures were grown by shaking at 30°C unless otherwise indicated. Where applicable, antibiotics were used at the following concentrations: streptomycin, 200 µg/mL; carbenicillin 100 µg/mL. 5-Bromo-4-chloro-3-indolyl-B-D-galactopyranoside (X-Gal; 120 µg/mL) was added to plates for blue-white screening, and sucrose (10%) was added to plates for counterselection against suicide vectors.

### Plasmid and strain construction

Plasmids were built using isothermal assembly [23] using primers listed in Table S1. Gene deletions were performed via homologous recombination using the suicide vector pCVD442 [24]. Chromosomal insertions were constructed using the vector pTD101, which inserts genes via double crossover into native *lacZ*.

All strains used in this study (Table S2) are derivatives of *V. cholerae* El Tor N16961 (WT). Plasmids were constructed and stored in DH5a, and conjugated into *V. cholerae* using *E. coli* SM10 or MFD donor strains. Liquid cultures of the donor strain were prepared in the appropriate antibiotic and mixed in equal ratio (10 µL + 10 µL) with the recipient strain on an LB plate. After overnight incubation at 37°C, cells were plated on LB containing streptomycin and carbenicillin to select for transconjugants. Colonies containing integration vectors were cured through two rounds of purification on salt-free sucrose agar containing streptomycin. Insertions and deletions were verified using PCR screening.

The *shyA::shyA*^Δ*lysM*^ strain was created by PCR amplifying sequences upstream and downstream of the annotated LysM domain using primers DLP213/214 (upstream homology) and DLP215/216 (downstream). Fragments were column-purified and cloned into Sma1-digested pCVD442 using isothermal assembly [23]. Deletion of *shyC* in this background was done using pCVDΔ*shyC* [9]. All strains were validated by whole-genome sequencing.

### sBADA and FM4-64 staining

Strains were inoculated from frozen stocks into 5 mL LB and incubated overnight at 30°C with shaking. After incubation, cells were diluted 1:1000 into LB or SF-LB. After 1.5 hours, sBADA was added to a final concentration of 125 µM. After 45 min shaking incubation at 37C, cells were harvested and imaged on an agarose pad (0.8% agarose in LB or SF-LB) containing FM4-64 (5 µg/mL). Cells were imaged using a Leica DMi8 inverted microscope, and resulting images were processed minimally using Leica LasX software.

### Time lapse microscopy

Cells were imaged under phase contrast on an agarose pad (0.8% agarose in LB or SF-LB) using a Leica DMi8 inverted microscope with frames taken every 5 min. Stage temperature was set to 37C using a PECON TempController 2000-1.

### Quantitative image analysis

Time lapse images were analyzed using SuperSegger [20] and the resulting values were graphed using custom R scripts.

### Growth curve analysis

Strains were grown overnight in LB and diluted 1:1000 into LB or SF-LB in a 100-well honeycomb plate. The growth of each well was monitored by optical density at 600 nm (OD600) on a Bioscreen C plate reader (Growth Curves America).

### Time-dependent killing assays

Cells were inoculated from frozen stocks into 5 mL LB and incubated overnight at 30C with shaking. After incubation, cells were diluted 1:100 into LB or SF-LB containing 100 ug/mL Penicillin G. Survival was measured by spot plating and calculating CFU/mL at indicated times.

## Acknowledgements

This work was supported by NIH-NIAID grants R01AI143704 and R01GM130971 to TD.

