## Supplemental Figure 2 for "Differential endopeptidase requirements during adaptation to changing growth conditions in *Vibrio cholerae*"

### Figure 2 Supplement

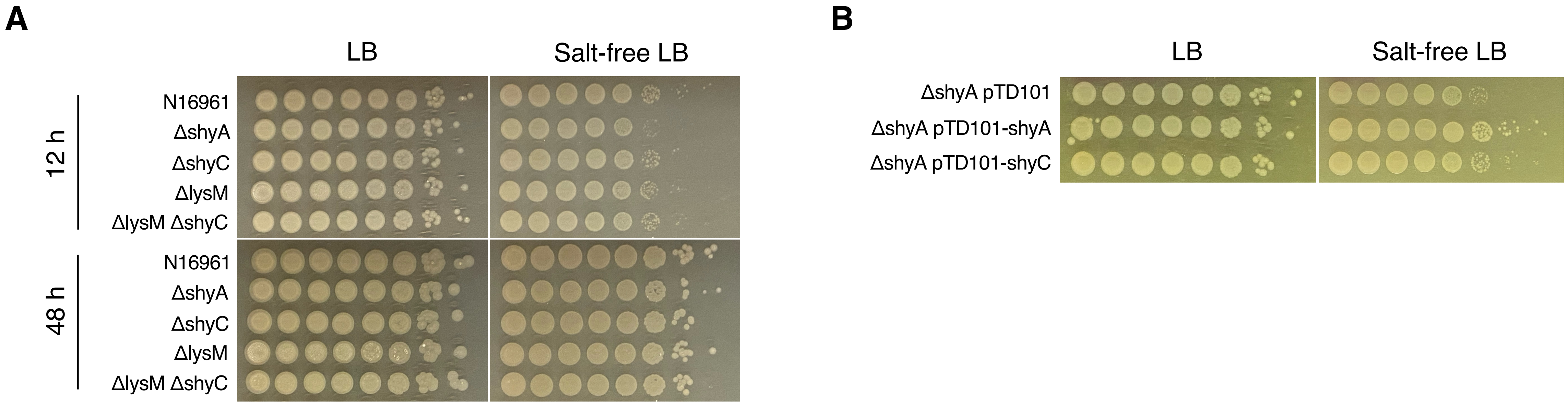


**Figure 4.** ShyC is not required for normal growth on low osmolarity medium and cannot fully rescue cells missing ShyA. **(A)** Spot dilutions of endopeptidase mutants were incubated at 37C on LB or salt-free LB and imaged after 12 hours and 48 hours. **(B)** Spot dilutions were incubated at 30C for 16 hours.
